# Identification of MIRNA-MRNA-HUB Gene Network Prediction in Squamous Cell Lung Carcinoma Via Starbase Server and Insilico Analysis of Hub Genes

**DOI:** 10.1101/2021.08.22.457301

**Authors:** Ashwini Kalaiselven, Venkataramanan Swaminathan

## Abstract

Malaysian has suffered from lung cancer, accounting for around 10% of all cancers reported in 2021. Prioritization of early diagnosis and natural treatments of lung cancer patients is crucial to improve their survival rate.The aim of this study is to identify the miRNA-mRNA and potential hub gene corresponding to lung carcinoma and natural drugs candidates to inhibit progression of lung cancer via insilico approaches.GSE176348,GSE85841 and GSE164750 gene expression profiles were retrieved from NCBI and GEO2R analysis was carried out accordingly to principal standard p<0.05 and logFC>1 to identify up-regulated and down-regulated genes from the datasets. Protein-protein interaction analysis of up-regulated and down-regulated genes was performed in Cytoscape software and top 10 hub genes were observed by cytohubba tools plugin in Cytoscape.Next, Biological Process (BP), cellular component (CC), molecular function (MF) of top 10 up regulated and down regulated hub genes were observed via DAVID server. The survival curves of top 10 up and down regulated hub genes were constructed by the Kaplan-Meier plotter and the expression level of hub genes in lung cancer tissues and normal tissues analyzed via GEPIA2 online database. IMPPAT: Indian Medicinal Plants, Phytochemistry And Therapeutics database was utilized for selection of potential natural drug candidates for lung cancer and ADME test was carried out on the selected drug compounds.To identify the most promising drug candidates,the hub genes were docked with respective natural drugs candidates via Swiss Dock online tool.Finally, The Encyclopedia of RNA Interactomes (ENCORI) database was used to study the MIRNA-MRNA network of targeted hub genes.In summary,this study is helpful in discovery of miRNAs-mRNA and analysis of natural drug candidates to inhibit lung carcinoma.

## 1) INTRODUCTION

Cancer is a disease phenomenon in which body cells grow out of control and spread to other parts of the body. Cancer is caused by changes to DNA where most cancer-causing DNA changes occur in sections of DNA called genes. These changes are also called genetic changes which can contributing in tumour formation. A tumour might be cancerous (malignant) or noncancerous (benign).

Lung carcinoma is a type of cancer that begins in the lungs and has no symptoms or indicators in its early stages. Symptoms and indications of lung cancer usually appear after the disease has progressed. The majority of lung cancers do not cause symptoms until the disease has advanced, and the symptoms can vary from person to person. Lung cancer symptoms include pain in the chest, back, or shoulders that gets worse with coughing, deep breathing, shortness of breath, unexplained weight loss, exhaustion, lack of appetite, and hoarseness or wheezing.

Chemotherapy is one way used to treat cancer, however because to the lack of drug selectivity, a substantial percentage of healthy cells are destroyed along with malignant cells. The most difficulty in cancer treatment is destroying tumour cells in the presence of healthy cells without harming the healthy cells. To create anti-cancer medications from natural resources such as plants, cytotoxic chemicals must be tested and crude plant extracts must be studied.(Kooti et al. 2017)

The aim of this study were to study the PPI network of lung cancer and to determine the cellular composition, biological processes and molecular functions of gene expression. On the other hand, the discovery of central gene biomarkers helps in studying the overall survival and the onset of lung cancer expression. Accordingly, bioinformatics tools are important for high-throughput sequencing, which will improve analysis of early diagnosis, staging, and prognosis. We determine the suitable natural drug target that can inhibit the targeted lung carcinoma gene.

## 2) MATERIALS & METHODS

### 3.1 DATA COLLECTION

#### 3.1.1 Retrieval and selection of data from GEO datasets from NCBI

To carry out this study, the data was extracted from the NCBI website (https://www.ncbi.nlm.nih.gov/gds). The data extracted was about lung carcinoma under. From 818 search results, three different lung carcinoma GEO accession ID’s GSE176348,GSE85841 and GSE164750 were chosen for this study.Each accessions contains 24 samples,16 samples and 27 samples respectively.

#### 3.1.2 Data Processing and DEGs Selection

The chosen three accession ID’s were run through GEO2R analysis which was available in NCBI. The DEGs screened accordingly principal standard p<0.05 and logFC>1 from the dataset.The upregulated and downregulated DEGs are considered as logFC ≥ 1 and log FC ≤ 1.This step applied for these three accession ID GSE176348,GSE85841and GSE164750 respectively.

#### 3.1.3 Determination of Coexpression Modules

All the DEG genes, both upregulated and downregulated genes of this three accession ID’s GSE176348,GSE85841and GSE164750 were grouped.The intersection nodes of GSE176348,GSE85841and GSE164750 accesion ID DEG genes were identified and displayed in a venn diagram via online venn diagram tools (http://bioinfogp.cnb.csic.es/tools/venny/) as shown in figure 2.

**FIGURE 1:**
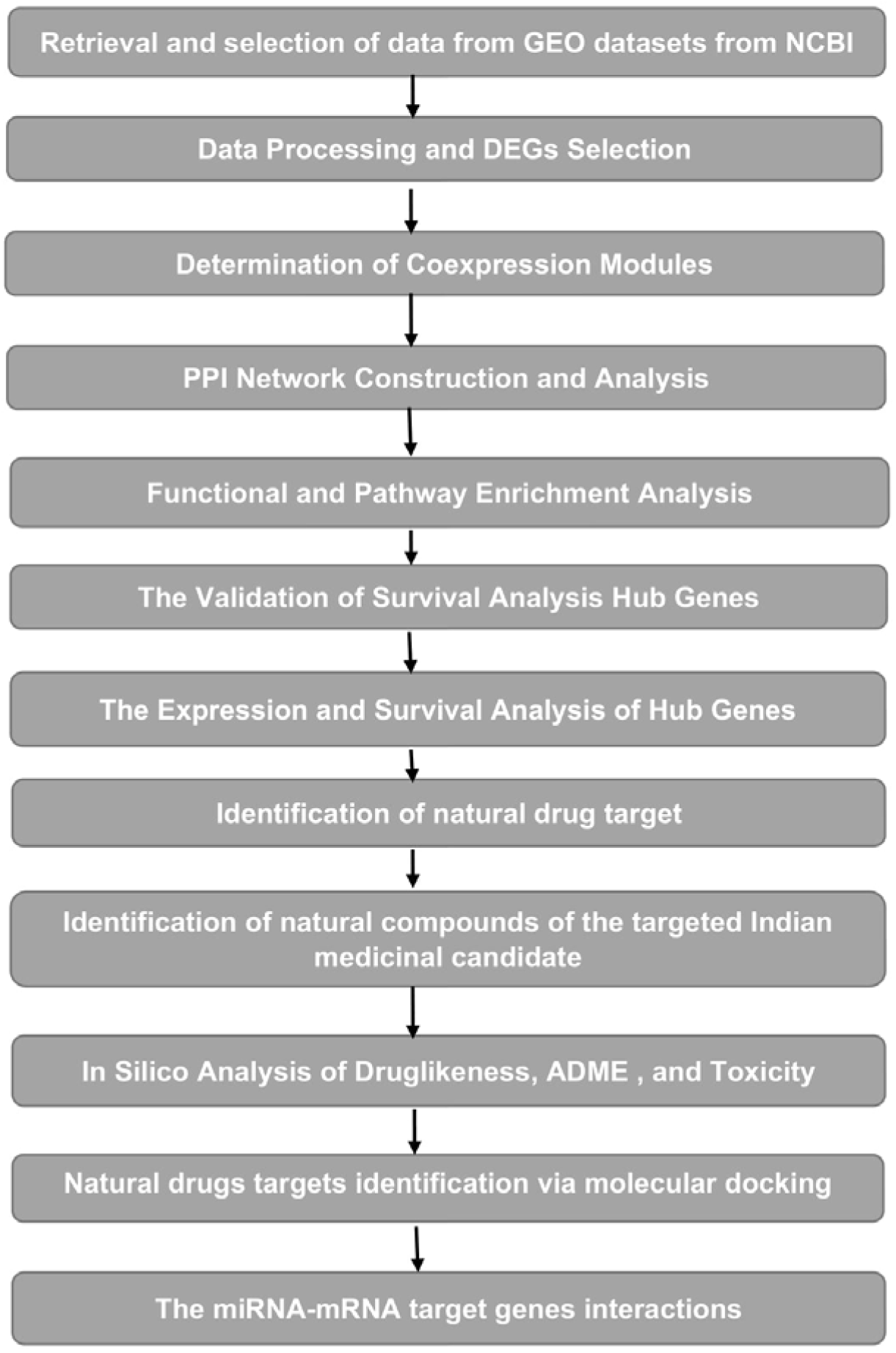
FLOWCHART OF METHODOLOGY.

**FIGURE 2:**
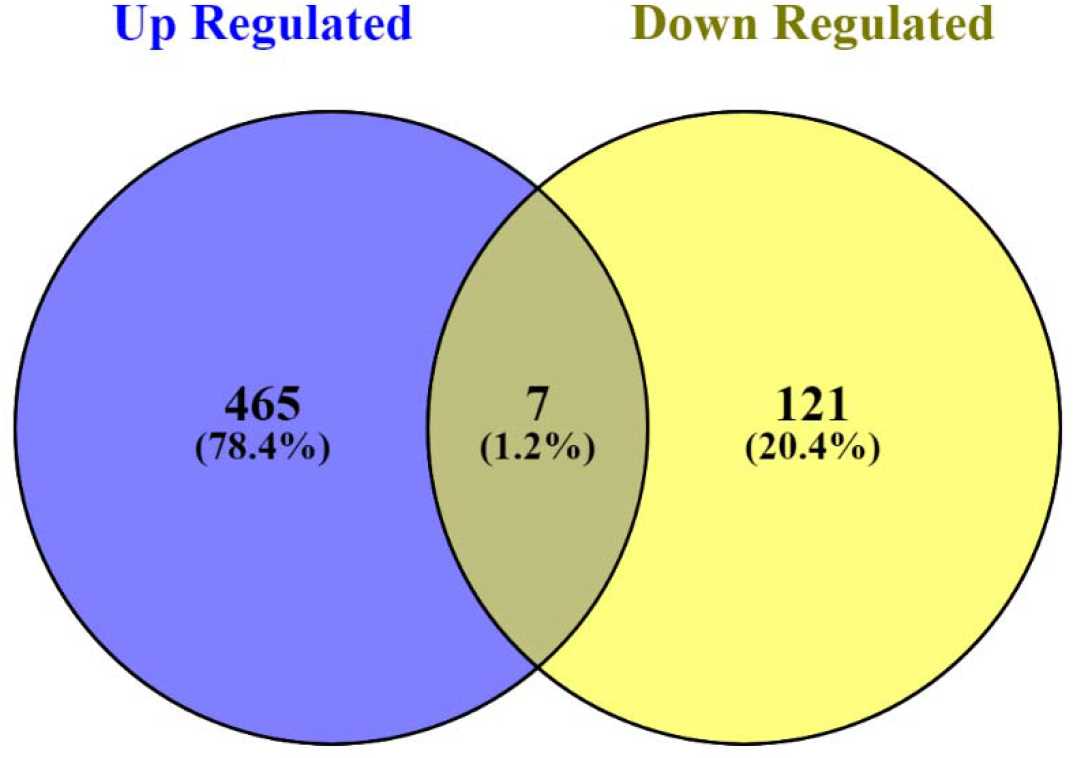
Venn diagram shows the total number of upregulated genes, downregulated genes and their intersection genes

### 3.2 DATA ANALYSIS

#### 3.2.1 PPI Network Construction and Analysis

Protein-protein interaction analysis of up-regulated genes and down-regulated genes for these three accession ID was performed in cytoscape software. Throughout this research, the protein-protein interaction setting was set with the confidence (score) cutoff to 0.4 and their respective maximum additional interactors to zero to import the network.Top 10 hub genes for both upregulated and downregulated genes were identified via cytohubba tools plugin in cytoscape and tabulated (Table 1 and Table 2).Protein-protein interactions diagrams for both upregulated and downregulated genes list were constructed.

**TABLE 1:**
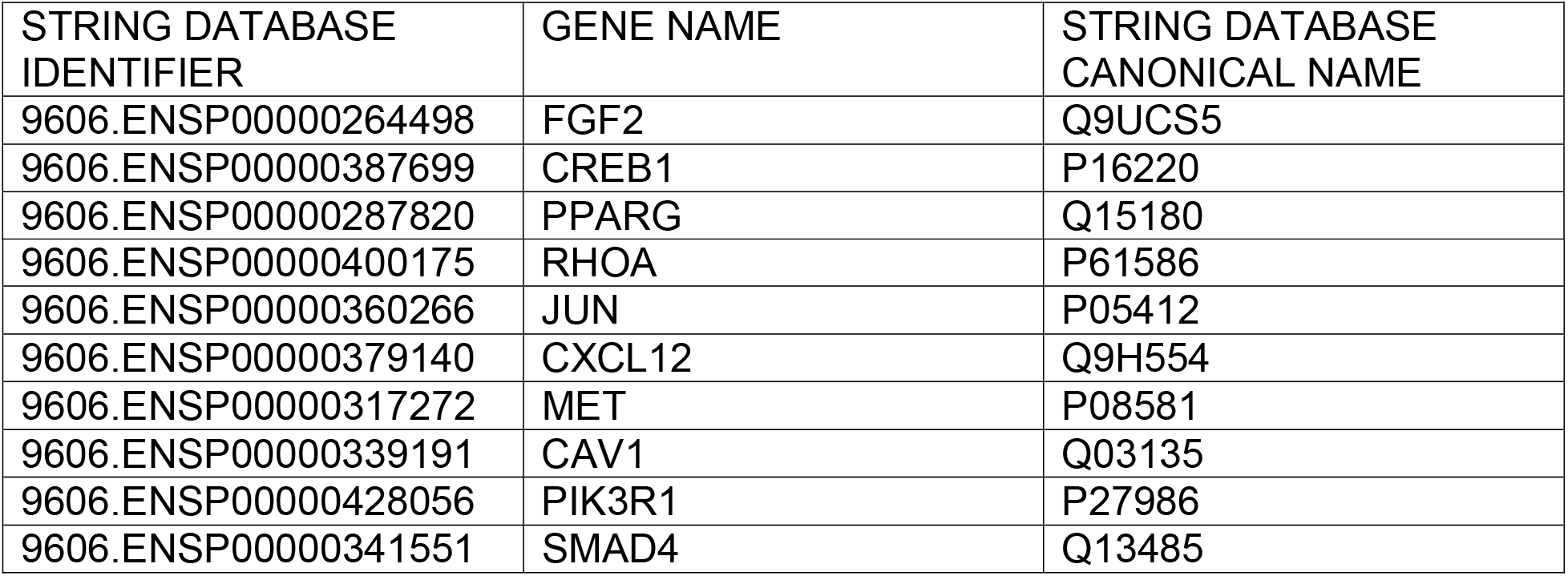
List of top 10 hub genes of up-regulated genes with their string database identifiers and canonical names of lung carcinoma

**TABLE 2:** List of top 10 hub genes of down-regulated genes with their string database identifiers ad canonical names of lung carcinoma

| STRING DATABASE IDENTIFIER | GENE NAME | STRING DATABASE CANONICAL NAME |
| --- | --- | --- |
| 9606.ENSP00000381072 | SERPINB8 | P50452 |
| 9606.ENSP00000378517 | SPP1 | P10451 |
| 9606.ENSP00000370201 | TAF9 | Q9Y3D8 |
| 9606.ENSP00000370193 | AK6 | Q16594 |
| 9606.ENSP00000366563 | PIK3CD | O00329 |
| 9606.ENSP00000364037 | TEX10 | Q9NXF1 |
| 9606.ENSP00000348168 | GTF2E2 | P29084 |
| 9606.ENSP00000320084 | CD276 | Q5ZPR3 |
| 9606.ENSP00000282397 | FLT1 | P17948 |
| 9606.ENSP00000244007 | PLCG1 | P19174 |

#### 3.2.2 Functional and Pathway Enrichment Analysis

DAVID servers (https://david.ncifcrf.gov/tools.jsp) associated Gene Ontology function and KEGG pathway enrichment with the functional annotation.The results of Biological Process (BP), cellular component (CC), molecular function (MF) were observed and determined. DEGs of the enrichment pathway, molecular, and biological function GO with adjusted significant value (p-values 0.05 and FDR0.05) were investigated.The results were tabulated in table 3 and 4.

**TABLE 3:** Functional enrichment analysis of Up regulated genes of lung carcinoma

| Category | Term |
| --- | --- |
| Biological Processes | Host-virus interaction |
| Molecular Function | Activator<br>Growth factor |
| Cellular Component | Secreted<br>Nucleus<br>Membrane<br>Cytoplasm<br>Golgi apparatus<br>Cytoskeleton<br>Cell projection |

**TABLE 4:** Functional enrichment analysis of Down regulated genes of lung carcinoma

| Category | Term |
| --- | --- |
| Biological Processes | Chemotaxis, Host-virus interaction, Transcription, Transcription regulation, Biological rhythms, Differentiation, Angiogenesis, Stress response, Protein transport, Cell cycle, Cell division |
| Molecular Function | Cytokine, Growth factor, Activator, DNA-binding, Kinase, Receptor, Transferase, Tyrosine-protein kinase, Developmental protein, Growth factor, Heparin-binding, Mitogen, Developmental protein, |
| Cellular Component | Secreted, Nucleus, Golgi apparatus, Membrane, Cell membrane, Cell projection, |

#### 3.2.3 The Validation of Survival Analysis Hub Genes

The Kaplan-Meier plotter (http://kmplot.com/analysis/index.php?p=service&cancer=lung) was accomplished on the top 10 up regulated and down regulated genes. The Kaplan Meier plotter is capable to assess the correlation between the expression of 30000 genes (mRNA, miRNA, protein) and survival in more than 25000 samples from 21 tumor types including breast, ovarian, lung and gastric cancer.The Primary purpose of this method is to investigate meta-analysis based discovery and validation of survival biomarkers.The results were tabulated in figure 6 and figure 7.

**FIGURE 3:**
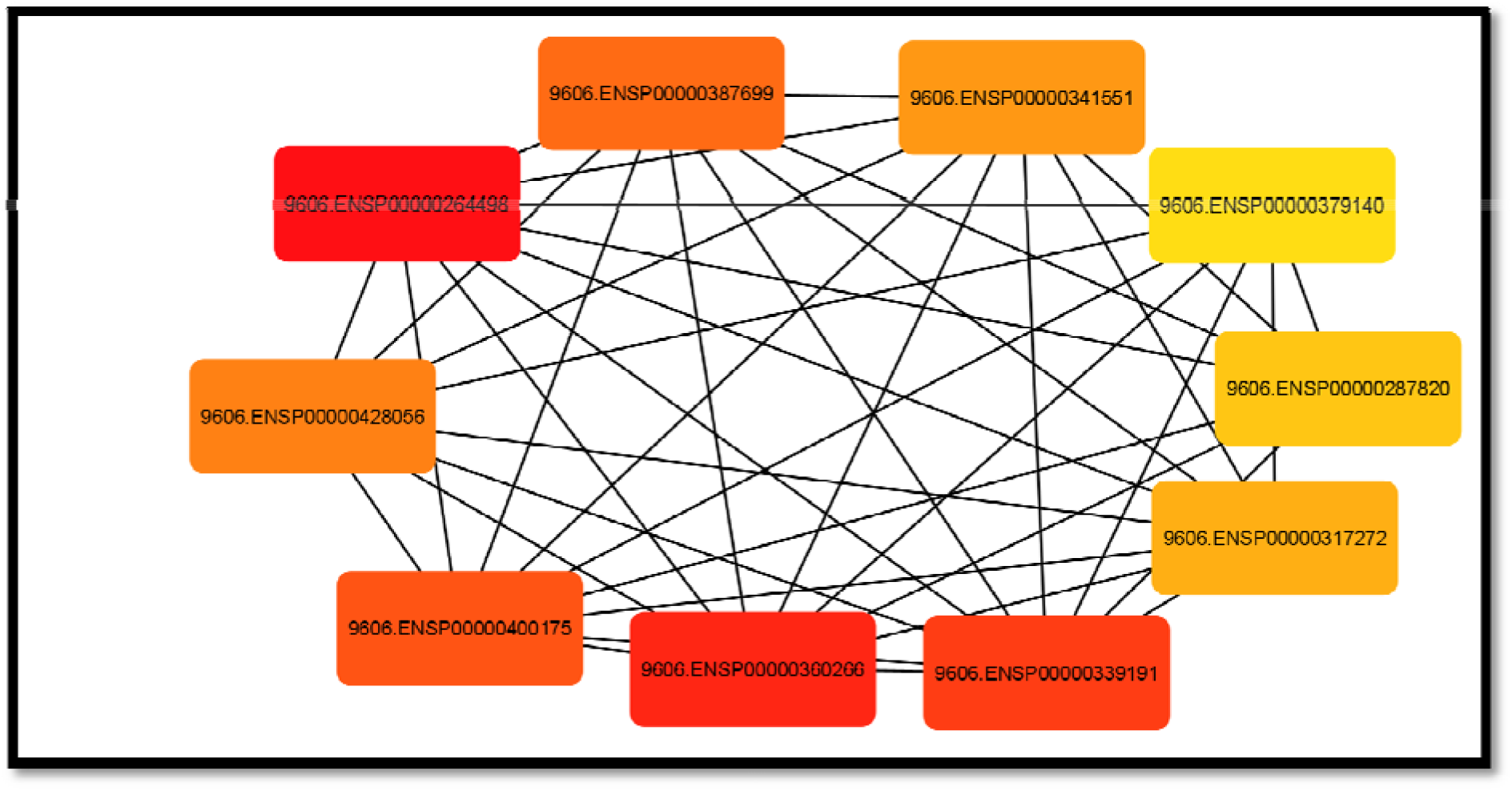
Protein-protein interaction diagram of top 10 hub genes of up regulated genes of lung carcinoma

**FIGURE 4:**
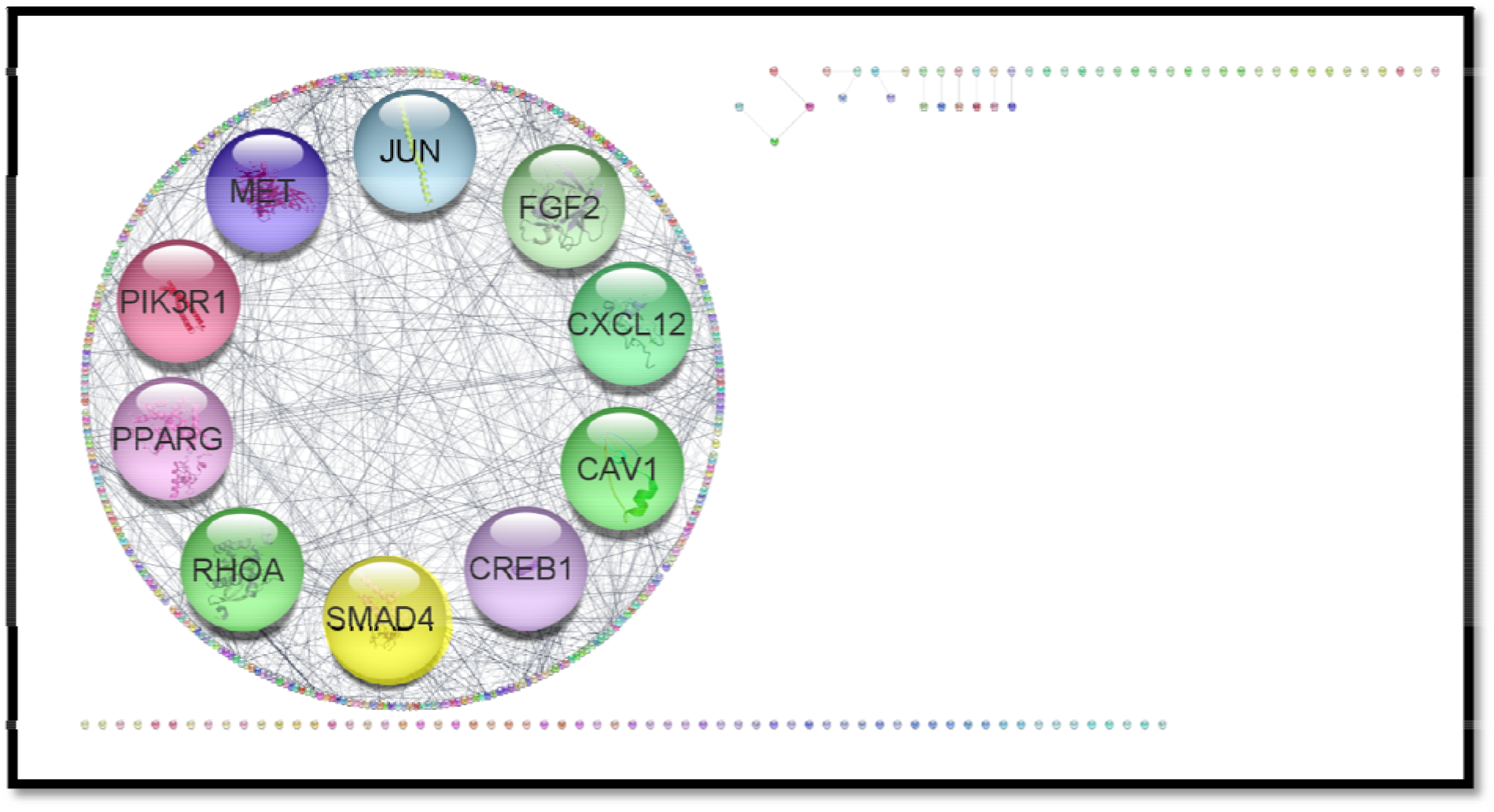
Protein-protein interaction diagram of up regulate lung carcinoma genes.

**FIGURE 5:**
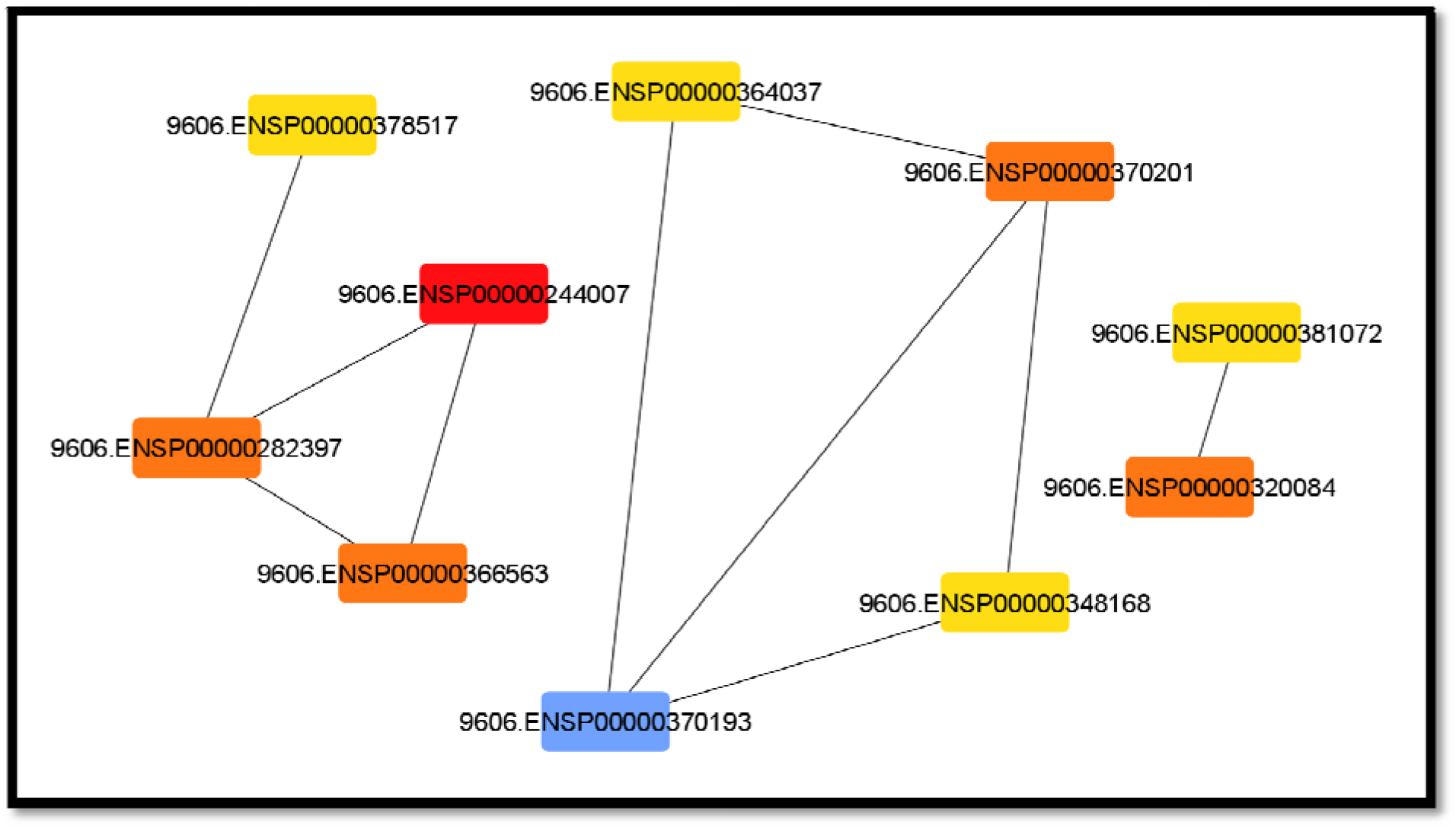
Protein-protein interaction diagram of top 10 hub genes of down regulated genes of genes of lung carcinoma

**FIGURE 6:**
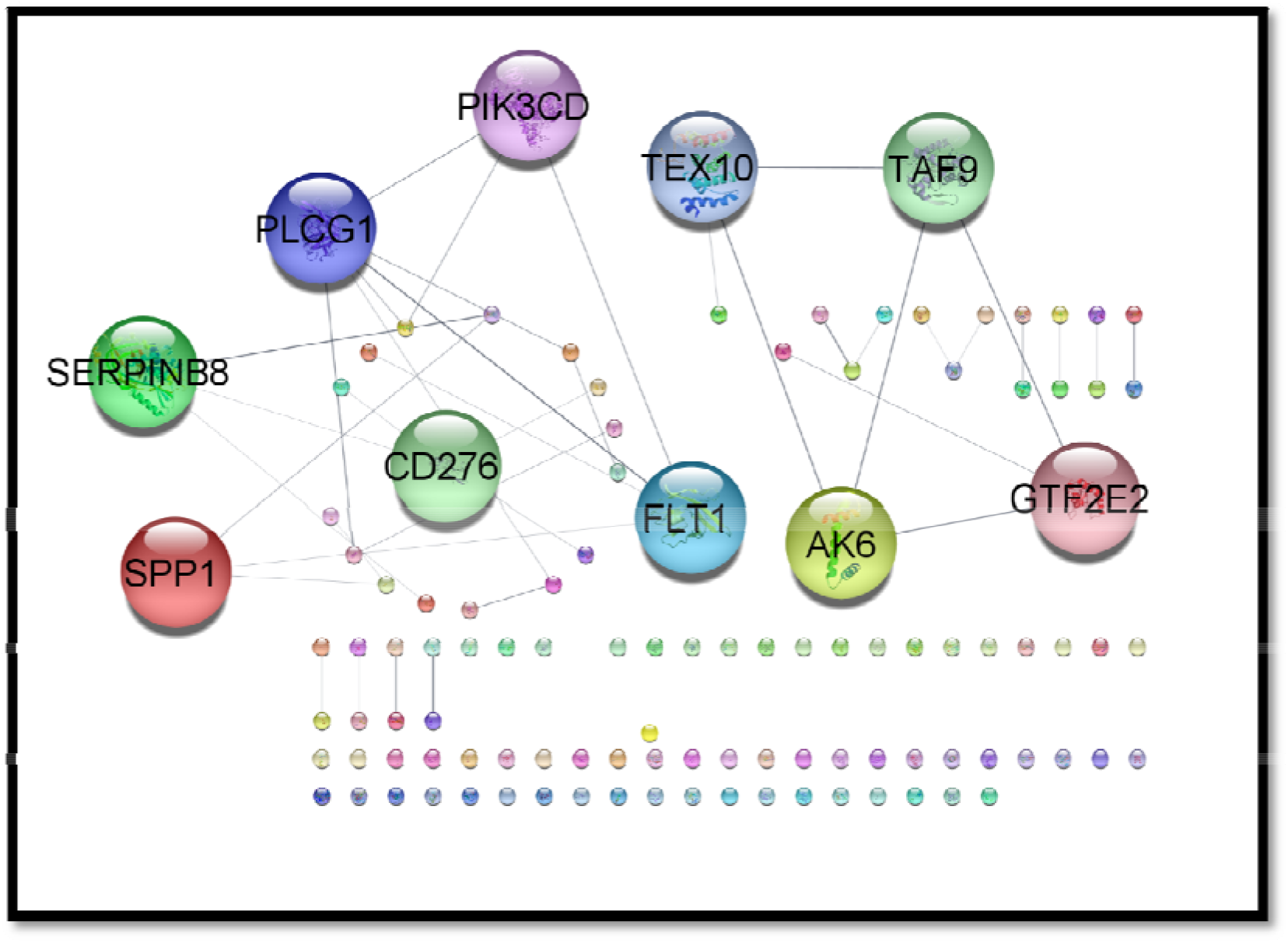
Protein-protein interaction diagram of down regulated genes of lung carcinoma

**FIGURE 7:**
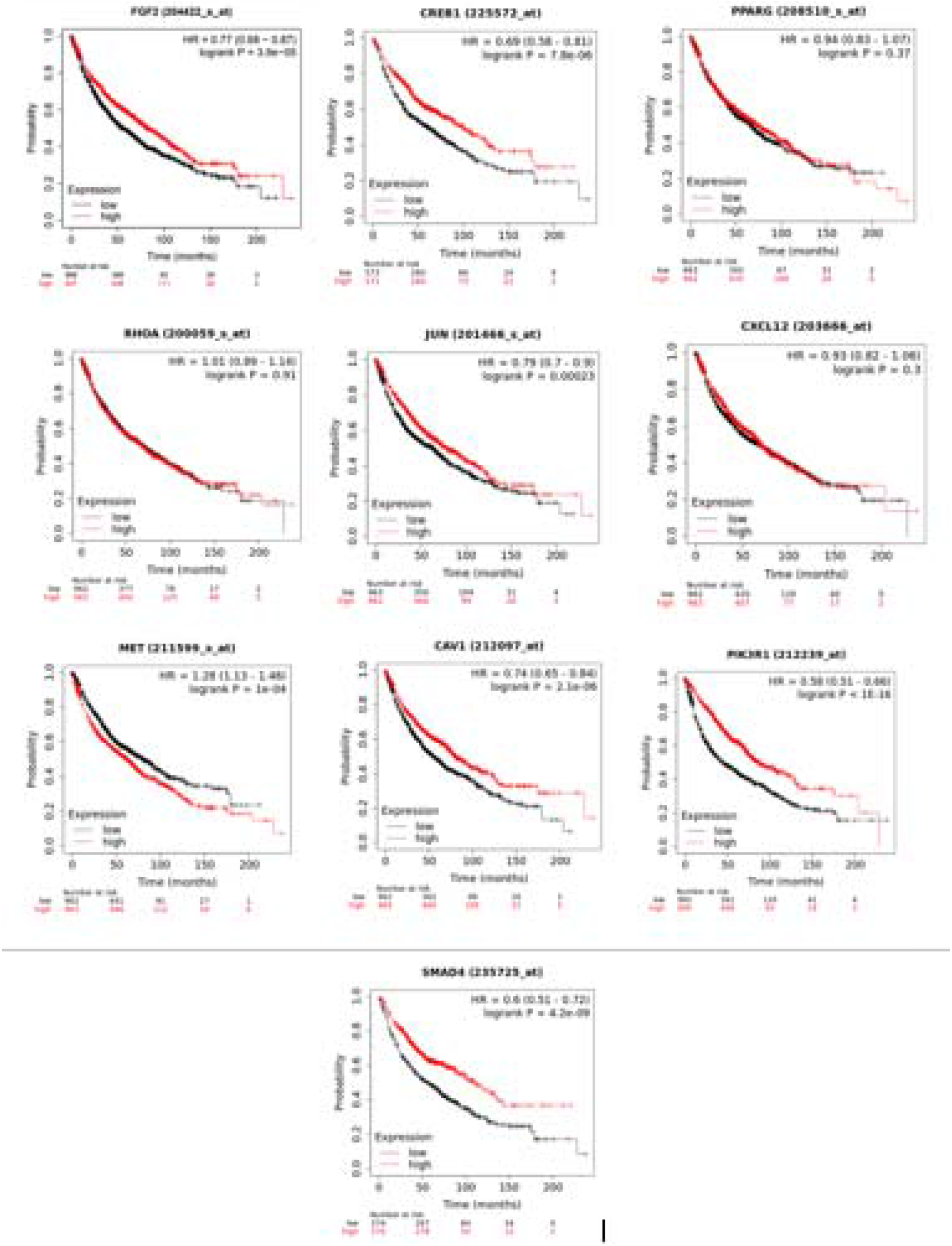
Survival Analysis Of Top 10 Up Regulated Genes (FGF2,CREB1,PPARG,RHOA,JUN,CXCL12,MET,CAV1,PIK3R1,SMAD4) of lung carcinoma

#### 3.2.4 The Expression and Survival Analysis of Hub Genes

Gene Expression Profiling Interactive Analysis (GEPIA) (http://gepia.cancer-pku.cn) was utilized to validate the expression of hub genes in TCGA-STAD tumor samples. To further analyze the effects of the hub genes on GAC patient prognosis survival, the GEPIA website was selected to draw overall survival (OS) curves. Meanwhile, we selected quartile as the group cutoff. Genes that significantly affect the prognosis survival of GAC patients were further analyzed for correlation with each other in the GEPIA website. The protein levels of these genes were obtained from the Human Protein Atlas (https://www.proteinatlas.org/, accessed on 26 January 2021). The results were tabulated in figure 8 and figure 9.

**Figure 8:**
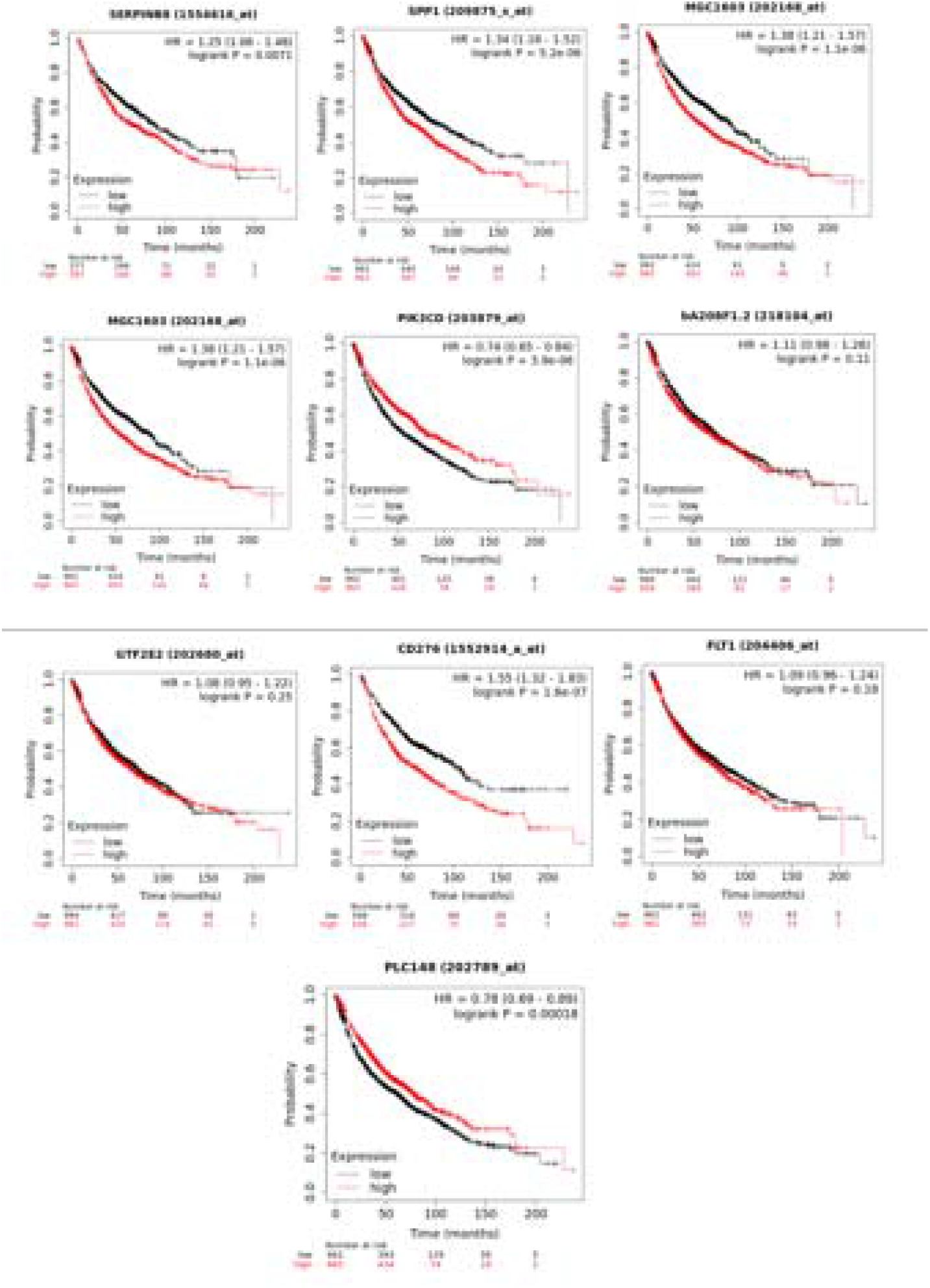
Survival Analysis Of Top 10 Down Regulated Genes (SERPINB8,SPP1,TAF9,AK6,PIK3CD,TEX10,GTF2E2,CD276,FLT1,PLCG1) of lung carcinoma

**FIGURE 9:**
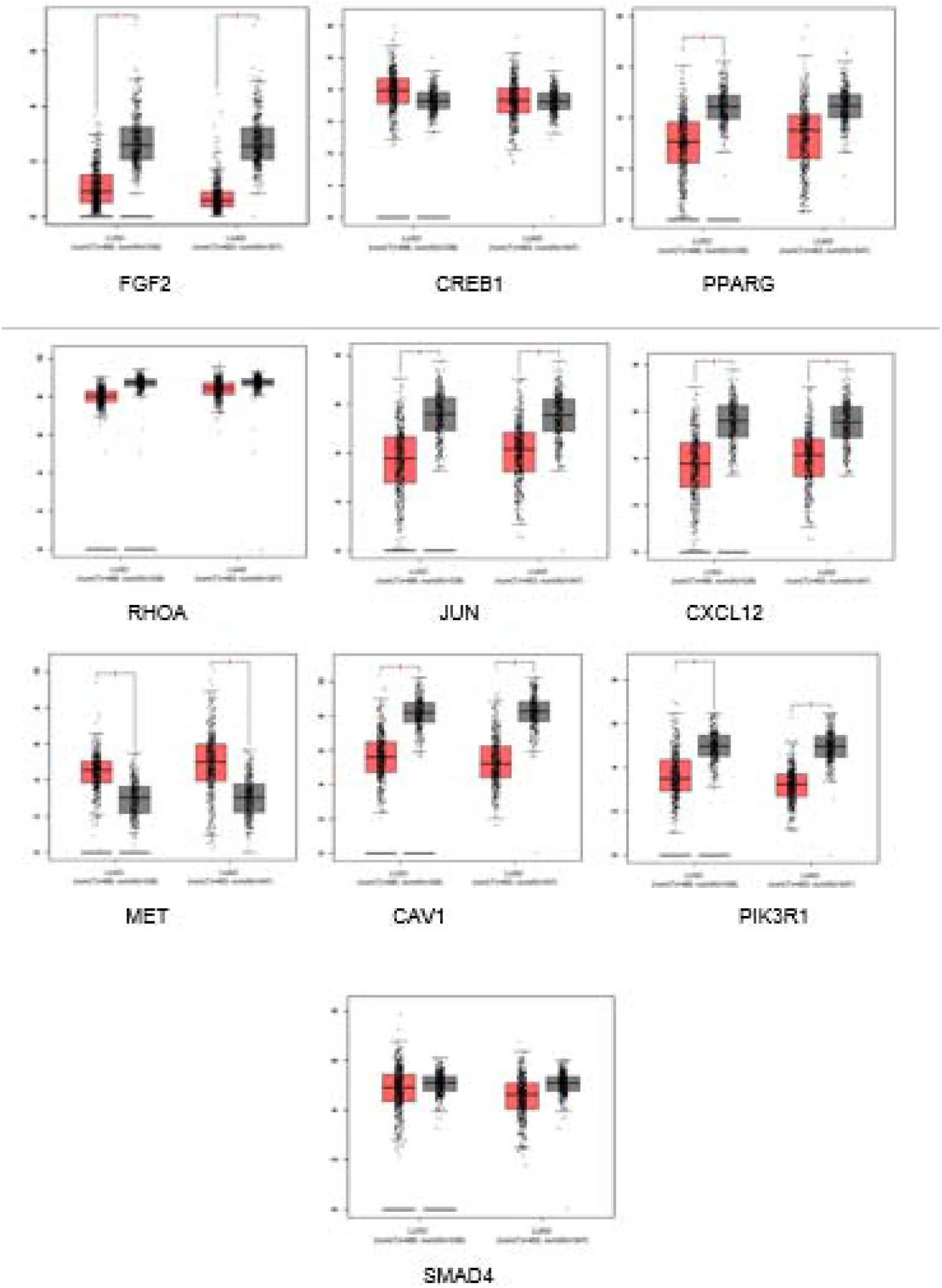
The expression level Of Top 10 Down Regulated Genes (FGF2,CREB1,PPARG,RHOA,JUN,CXCL12,MET,CAV1,PIK3R1,SMAD4) of lung carcinoma

#### 3.2.5 Identification of natural drug target

Indian medicinal candidate that can assist in treating lung carcinoma was identified by selecting lung cancer and lung carcinoma in therapeutic use filter through IMPPAT: Indian Medicinal Plants, Phytochemistry And Therapeutics database (https://cb.imsc.res.in/imppat/home). Allium sativum, Arnebia euchroma and Nigella sativa was selected as the Indian medicinal candidate for this study.

#### 3.2.6 Identification of natural compounds of the targeted Indian medicinal candidate

Allicin, acetylshikonin and thymoquinone was identified as the natural compounds of Allium sativum, Arnebia euchroma and Nigella sativa respectively. Allicin, acetylshikonin and thymoquinone are the core molecules present in the the Indian medicinal candidates that we have targeted for this study since previous docking studies was successful against docking.The results are tabulated in table 5.

**TABLE 5:** Tabulation of Indian drug target and the targeted compound.

| Indian medicinal candidate | The target compound |
| --- | --- |
| Allium sativum | Allicin |
| Arnebia euchroma | Acetylshikonin |
| Nigella sativa | Thymoquinone |

#### 3.2.7 In Silico Analysis of Druglikeness, ADME, and Toxicity analysis

The Druglikeness, ADME, and Toxicity analysis was carried out on Allicin, Acetylshikonin and thymoquinone compounds via (https://preadmet.qsarhub.com/toxicity/) software.The purpose of this analysis is to understand the safety and efficacy of a drug candidate in treating lung carcinoma.The results are tabulated in table 6.

**TABLE 6:** Tabulation and Comparison of Druglikeness, ADME, and Toxicity analysis of Ligands

| Description | Allicin | Acetylshikonin | Thymoquinone |
| --- | --- | --- | --- |
| Lipinski's "Rule-of-Five," | Suitable | Suitable | Suitable |
| BBB | 1.22158 | 0.0662739 | 1.78938 |
| Caco2 | 21.722 | 17.3775 | 23.037 |
| CYP_3A4_INHIBITION | NON | INHIBITOR | NON |
| HIA | 98.312414 | 91.239925 | 99.286337 |
| MDCK | 27.7678 | 29.017 | 61.0892 |
| PPB | 4.346351 | 91 | 100 |

#### 3.2.8 Natural drugs targets identification via molecular docking

PDB files of top 10 hub genes of both up regulated and down regulated are retrieved from Protein Data Bank (PDB) (https://www.rcsb.org/). Each hub genes PDB files were submitted one at a time to SwissDock (http://www.swissdock.ch/docking) to conduct molecular docking with respective natural drugs candidates selected which were identified from IMPPAT: Indian Medicinal Plants, Phytochemistry And Therapeutics database.The molecular docking results were tabulated in table 7 and 8. Drugs candidates that show high energy value will be selected for natural drug substitutes identification analysis.

**TABLE 7:** SwissDock molecular docking results of top 10 up regulated genes with three targeted compounds respectively. PIK3R1 shows no results in the compound Acetylshikonin due to error in molecular docking analysis due to unavailability of drugs structures.

|  | Allicin |  | Acetylshikonin |  | Thymoquinone |  |
| --- | --- | --- | --- | --- | --- | --- |
| GENE NAME | FullFitness (kcal/mol) | Estimated $\Delta G$(kcal/mol) | FullFitness (kcal/mol) | Estimated $\Delta G$(kcal/mol) | FullFitness (kcal/mol) | Estimated $\Delta G$(kcal/mol) |
| FGF2 | -2427.77 | -6.09 | -2462.82 | -8.07 | -2425.97 | -6.34 |
| CREB1 | -2192.06 | -6.29 | -2224.04 | -7.20 | -2188.54 | -6.25 |
| PPARG | -3412.27 | -6.71 | -3437.11 | -8.75 | -3407.21 | -6.56 |
| RHOA | -1266.63 | -6.23 | -1299.15 | -7.97 | -1264.44 | -6.14 |
| JUN | -2379.83 | -5.83 | -2410.18 | -6.65 | -2376.82 | -5.85 |
| CXCL1<br>2 | -767.60 | -6.05 | -798.33 | -8.01 | -764.35 | -6.16 |
| CAV1 | -1221.99 | -5.79 | -1255.60 | -8.26 | -1217.83 | -5.77 |
| PIK3R1 | -4703.08 | -6.63 | - | - | -4694.97 | -6.21 |
| SMAD4 | -2572.90 | -6.10 | -2607.36 | -7.15 | -2573.29 | -6.52 |

#### 3.2.9 Identification of miRNA-mRNA-hub gene network prediction via starbase server

The Encyclopedia of RNA Interactomes (ENCORI) is an open-source platform mainly focusing on miRNA-target interactions (http://starbase.sysu.edu.cn), miRNA-mRNA sub-module was selected under “miRNA-Target” module, with the specifications as clade (mammal), genome (human) and assembly (hg19) in starbase server.Each of the top 10 up regulated and down regulated hub genes were deposited one at a time and the results were tabulated in table 9 and 10. (Zhou et al. 2020)

**TABLE 9:**
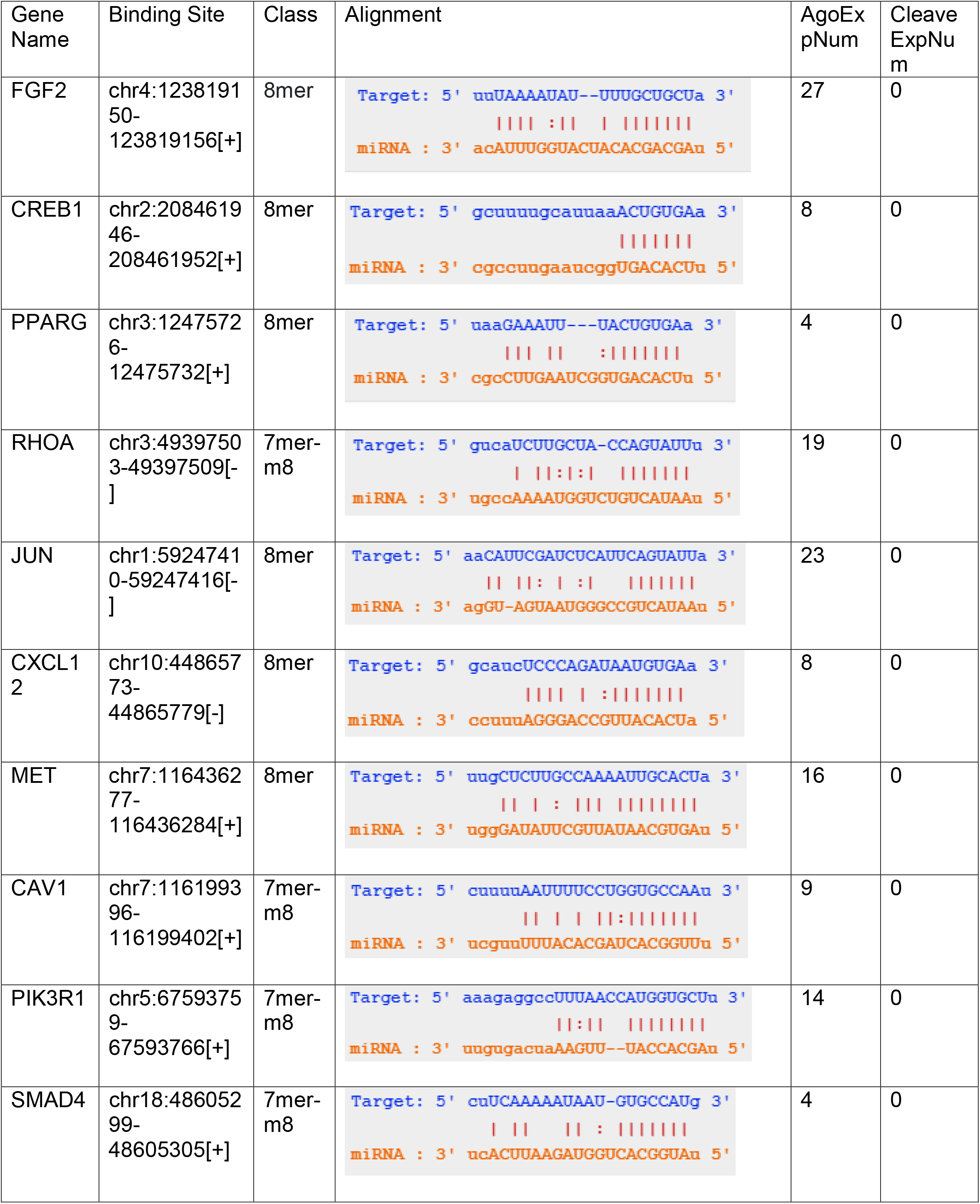
The miRNA-mRNA interactions of top 10 up regulated hub genes of lung carcinoma.

**TABLE 10:**
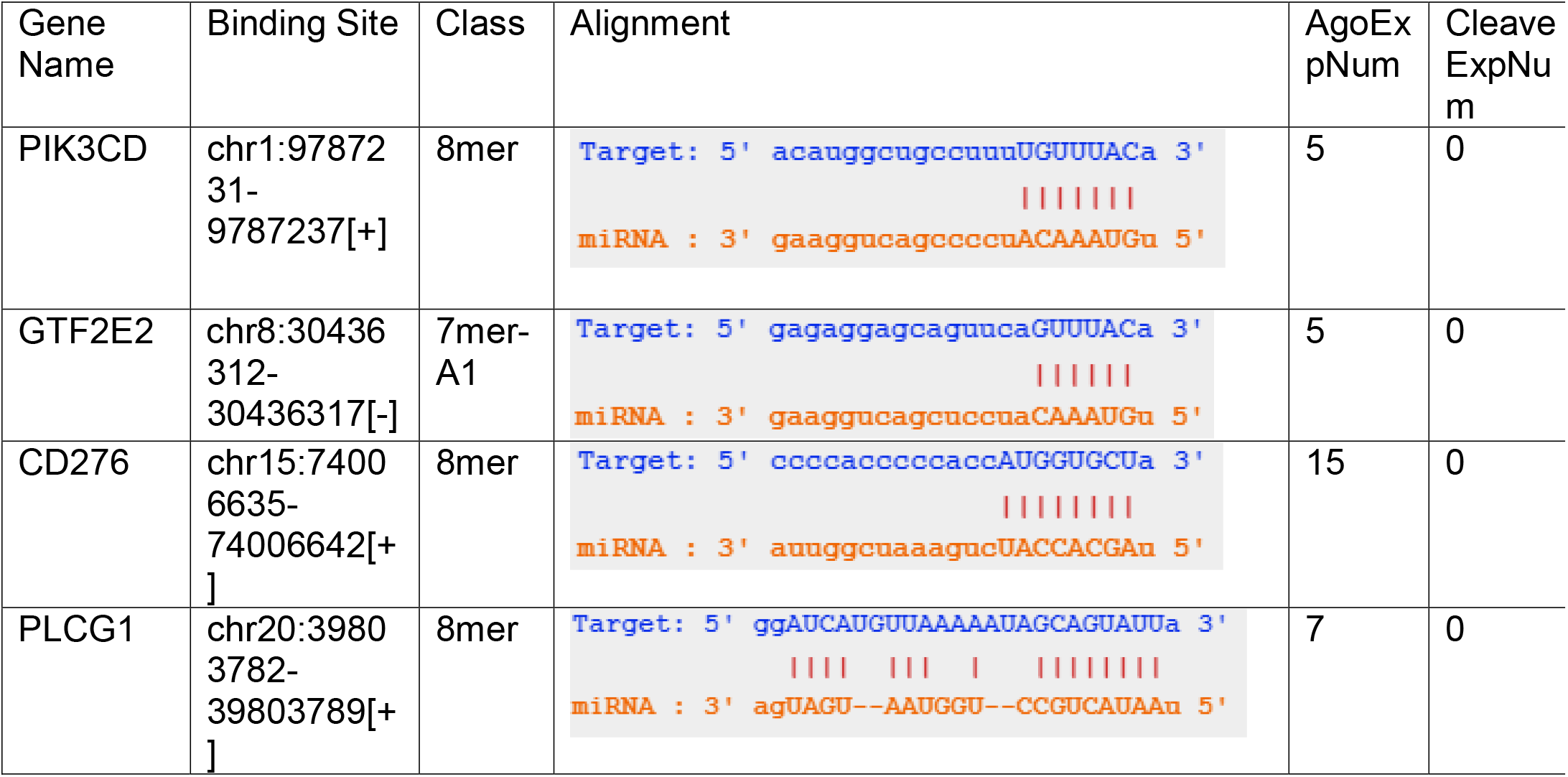
The miRNA-mRNA interactions of top 10 down regulated hub genes of lung carcinoma.

## 3) RESULTS

From this study, identified top 10 hub genes of lung carcinoma up-regulated genes are FGF2,CREB1,PPARG,RHOA,JUN,CXCL12,MET,CAV1,PIK3R1,SMAD4 while top 10 hub genes for lung carcinoma down-regulated genes are SERPINB8, SPP1, TAF9, AK6, PIK3CD, TEX10, GTF2E2, CD276, FLT1, PLCG1 which are shown table 1 and table 2. The protein-protein interaction of up regulated and down regulated hub genes of lung carcinoma was studied and displayed in figures 3, 4, 5 and 6. The Biological Process (BP), cellular component (CC), molecular function (MF) of top 10 up regulated and down regulated hub genes of lung carcinoma was tabulated in table 3 and 4.

Survival Analysis and The expression level of top 10 up regulated and down regulated hub genes of lung carcinoma was analyzed and displayed in figure 7,8,9 and 10.In this study, Allium sativum,Arnebia euchroma and Nigella sativa were resulted to be the most promising drug target for lung carcinoma, the results of molecular docking of the compounds and drug targets are shown in table 7 and 6.The miRNA-mRNA interactions of top 10 up regulated and down regulated hub genes of lung carcinoma are tabulated in table 9 and 10.

**FIGURE 10:**
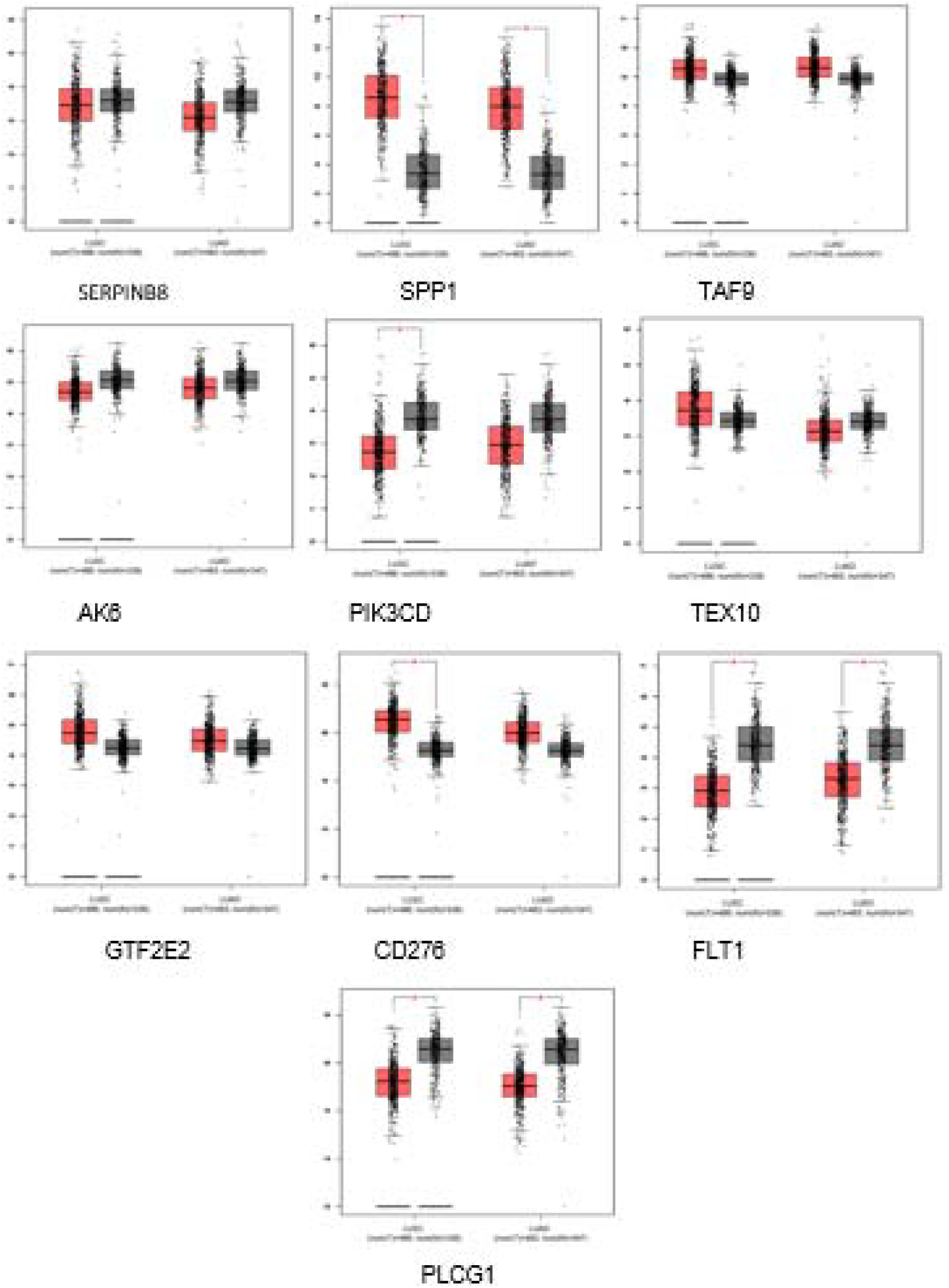
The expression level Of Top 10 Down Regulated Genes (SERPINB8, SPP1, TAF9,AK6,PIK3CD,TEX10,GTF2E2,CD276,FLT1,PLCG1) of lung carcinoma

## 4) DISCUSSION

The World Health Organization (WHO) defines lung adenocarcinoma categorization as a qualitative assessment of tumour differentiation based on both the extent to which the tumor’s architectural pattern resembles normal lung histology and cytologicatypia (Barletta, Yeap, and Chirieac 2010) When compared to other types of cancer patients, lung carcinoma patients have significant symptomatology. Because lung cancer patients have a low survival rate in the first year after diagnosis, concerns like palliation, death, and dying are frequently discussed at the time of diagnosis (Anon n.d.) The 5-year survival percentage for patients with Stage I (T1 or T2, N0, M0) disease is less than 70%. It’s possible that earlier identification of peripheral lung cancer will lead to a lower mortality rate (Anon n.d.)

Lung adenocarcinoma categorization is defined by the World Health Organization (WHO) as a qualitative assessment of tumour differentiation based on the extent to which the tumor’s architectural pattern mimics normal lung histology as well as cytologicatypia. Patients with lung carcinoma have more symptomatology than those with other forms of cancer. Because lung cancer patients have a poor first-year survival rate, issues such as palliation, death, and dying are commonly discussed at the moment of diagnosis. Patients with Stage I (T1 or T2, N0, M0) illness have a 5-year survival rate of less than 70%. It’s likely that detecting peripheral lung cancer earlier will result in a decreased mortality rate.

Therefore we have conducted this study prior in identifying indian medicinal drug targets that can inhibit against lung carcinoma through series of bioinformatics in-silico analysis.Throughout this study, three lung carcinoma accession ID’s were extracted from GEO Datasets NCBI,which are GSE176348,GSE85841 and GSE164750.Series of analysis were conducted on this ID’s.Each samples of respective accession ID’s, were grouped as tumorous and non-tumorous and ran through GEO2R analysis for the identification of their up regulated and down regulated gene sets.

Gene regulation is an important part of normal development. Genes are turned on and off in different patterns during development to make a brain cell look and act different from a liver cell or a muscle cell, for example. Gene regulation also allows cells to react quickly to changes in their environments.The identification of up regulated and down regulated genes is to get the over and low working genes than the optimum level. Gene expression is regulated to ensure that the correct proteins are made when and where they are needed. Regulation may occur at any point in the expression of a gene, from the start of the transcription phase of protein synthesis to the processing of a protein after synthesis occurs. All the up regulated and down regulated gene were grouped for further analysis.With the grouped up regulated and down regulated gene,we identify the intersection nodes via online Venn diagram tools.As the result, we had seven genes which was repeated in both up regulated and down regulated genes list,465 up regulated genes and 121 down regulated genes.

Next, protein-protein interaction analysis of up-regulated genes and down-regulated genes for these three accession ID was performed in cytoscape software.Protein-protein interaction plays key role in predicting the protein function of target protein and drug ability of molecules. The hub genes of both up-regulated and down-regulated genes lung carcinoma were identified and retrieved from CytoHubba tool in Cytoscape. From this study, identified top 10 hub genes of up-regulated genes of lung carcinoma are FGF2, CREB1, PPARG,RHOA,JUN,CXCL12,MET,CAV1,PIK3R1,SMAD4. Top 10 hub genes of lung carcinoma down-regulated genes are SERPINB8, SPP1, TAF9, AK6, PIK3CD, TEX10, GTF2E2,CD276,FLT1,PLCG1.

The top 10 hub genes of up-regulated and down-regulated genes of lung carcinoma was further utilized in functional annotation analysis conducted through DAVID servers to obtain the functional and pathway enrichment criteria of these potential hub genes corresponds to lung carcinoma. The results of Biological Process (BP), cellular component (CC), molecular function (MF) were determined.Based on the results, the hub genes of up-regulated genes were involved in host-virus interaction.This genes involves in viral invasion in the host,we also determined that these hub genes can be found in secreted,nucleus,membrane,Cytoplasm,Golgi apparatus,Cytoskeleton and Cell projection.Molecular Function describes the tasks performed by individual gene products like transcription factor and DNA helicase.(Dennis et al. 2003) The top 10 hub genes of up-regulated genes contributes in Activator and Growth factor as its molecular function.

The hub genes of down-regulated genes have involved in a few biological processes, those are Chemotaxis,Host-virus interaction,Transcription, Transcription regulation, Biological rhythms, Differentiation, Angiogenesis, Stress response, Protein transport, Cell cycle and Cell division.The hub genes of down-regulated genes mainly identified at Secreted, Nucleus, Golgi apparatus, Membrane, Cell membrane and Cell projection.They also acts as cytokine, Growth factor, Activator, DNA-binding, Kinase, Receptor, Transferase, Tyrosine-protein kinase, Developmental protein, Growth factor, Heparin-binding, Mitogen and Developmental protein.

Prognosis values of both up-regulated and down regulated hub genes of lung carcinoma were determined via KM plotter. The survival curves of 40 patients with lung cancer were examined using a specified parameter. Additionally, the GEPIA2 database was used to verify the expression levels of the 10 hub genes in tumor and normal tissues.The LUAD and LUSC of lung cancer with normal tissue samples are represented by the red and blue boxes.A median is the thick straight line in the middle. Each box’s lower and upper boundaries represented the first and third quartiles, respectively. The error bars in the bottom and top bars represented the expression data’s minimum and maximum values. The asterisk, which is statistically significant with each dot, denotes a tumour or normal tissues, respectively, for differential expression analysis.

Chemotherapy is one way used to treat cancer, however because to the lack of drug selectivity, a substantial percentage of healthy cells are destroyed along with malignant cells. The most difficulty in cancer treatment is destroying tumour cells in the presence of healthy cells without harming the healthy cells. To create anti-cancer medications from natural resources such as plants, cytotoxic chemicals must be tested and crude plant extracts must be studied.(Kooti et al. 2017).

For this study, potential natural medicinal candidates of lung cancer were referred and determined from IMPPAT database. Indian Medicinal Plants, Phytochemistry And Therapeutics (IMPPAT) is a curated database which has been constructed via literature mining followed by manual curation of information gathered from specialized books on traditional Indian medicine, more than 7000 abstracts of published research articles and other existing database resources. The schematic figure 1 gives an overview of the IMPPAT database construction pipeline.*Allium sativum*, Arnebia euchroma and Nigella sativa are proven to be effective in cancer treatments based on previous studies especially in treating lung cancer.

Allium sativum or practically called as garlic, considered as a conventional treatment for the majority of health-related issues.(Yeh et al. 2021) The major active element of Allium sativum is Allicin. Various research have shown that,Allicin have a clear and substantial biological effect on immune system enhancement, treatment of cardiovascular disorders, cancer, liver, and other areas. (Kooti et al. 2017). Arnebia euchroma is an endangered medicinal plant that grows natively in the Himalayas’ harsh cold and desert climates. (Xiong et al. 2009)This plant’s roots contain anti-inflammatory, antibacterial, and antipyretic qualities and have long been used in various medicinal needs.(Jain et al. 2021). Nigella sativa is called as black seed or black cumin, they contain thymoquinone,the main compound that has been used in traditional medicine to treat a variety of ailments, including cancer, asthma, hypertension, diabetes, inflammation, cough, bronchitis, headache, dermatitis, fever, dizziness, influenza and antioxidant protective effects.(Anon n.d.)

Due to their effectiveness in treating cancer, these 3 candidates will be utilized in further drug analysis via molecular docking approaches. Before proceeding to docking process, we have considered all compounds had undergo Absorption distribution metabolism excretion (ADME), toxicity and druglikeness testing.The reason for this process is to understand the safety and efficacy of a drug candidate, and are necessary for regulatory approval.

The ADME calculation gave details such as human intestinal absorption (%); in vitro Caco-2 cell permeability (nm/s), in vitro MDCK cell permeability (nm/s), in vitro skin permeability (logKpcm/h), in vitro plasma protein binding (%) and in vivo blood brain barrier penetration (c.blood/c.brain). The HIA (Human Intestinal Absorption) results demonstrate the best absorption of all the three natural drug compounds into human intestine. They showed moderate cellular permeability against Caco-2 cells.

Based on the ADME result on table 6, all the three compounds was resulted suitable in the most well-known rule/filter of drug-likeness which is Lipinski’s “Rule-of-Five,” which sets the boundaries of four simple molecular physicochemical criteria for orally active molecules. The results show that the compounds obey Lipinski rule of five and they can be strongly recommended as a drug candidate.(Anon n.d.).

BBB (Blood-Brain Barrier) penetration values help to know whether the compounds are able to pass across the blood-brain barrier or not. This parameter expresses the BBB penetration capacity and absorption rate of compound to CNS (Central Nervous System). Allicin,Acetylshikonin and Thymoquinone were observed to be having very moderate absorption to CNS.(G. Shibi et al. 2016)

All the selected lead compounds were found to be inhibitors of the protein CYP3A4, which demonstrates that they metabolize very easily. The PPB indicates the plasma protein binding of the drug and predicts its stay in the system and resultant clearance too. The PPB value of the compounds Acetylshikonin and Thymoquinone was highest among the selected natural drug candidates.(G. Shibi et al. 2016).

The natural drugs candidates were identified, molecular docking approaches were 10tilized to identify their energy value which is crucial in determining and selecting the good and more promising drug candidates. Drugs that are promising in inhibiting both upregulated and downregulated hub genes of lung carcinoma Allicin, Acetylshikonin and thymoquinone respectively.(Khan et al. 2011) Based on the docking results we can conclude that, Allicin is the best promising drug candidate to inhibit with the targeted genes, this is because the high score ranking compare to other compounds. Further study on the effectiveness of Allicin on these hub genes need to conducted to develop the potential drugs in treating lung carcinoma in the further.

Furthermore, MicroRNAs (miRNAs) are non-coding RNA that plays a significant role in tumor and its development by targeting genetic mutations or oncogenic genes.They usually regulate their target genes by binding to the complementary sequence at the 3’untranslated region.(Zhou et al. 2020) In this study, the top 10 up regulated and down regulated hub genes of lung carcinoma were analysed into ENCORI (RNA INTERACTIONS).

The binding sites of each up regulated hub genes was identified,FGF2 (CHR 4),CREB1(CHR 2), PPARG (CHR 3), RHOA (CHR 3),JUN (CHR 1),CXCL12 (CHR 3),MET(CHR 7),CAV1 (CHR 7), PIK3R1(CHR 5),SMAD4 (CHR 18). PPARG, RHOA and CXCL12 has the same binding sites which is at chromosome 3.The down regulated hub genes binds at PIK3CD (CHR 1), GTF2E2 (CHR 8), CD276 (CHR 15) and PLCG1 (CHR 20). MiRNAs detect their target mRNAs based on sequence complementarity and act on them to suppress protein translation through mRNA destruction. MicroRNA’s misregulation causes tumours, and they appear as oncogenes or tumour suppressors.(Peng and Croce 2016)

## 5) CONCLUSION

Throughout this study, we discovered up regulated and down regulated genes which are corresponded to lung carcinoma via differential gene expression analysis (DEG) via GEO2R analysis. The hub genes of both up-regulated and down-regulated genes lung carcinoma were identified and retrieved from CytoHubba tool in Cytoscape. From this study, identified top 10 hub genes of up-regulated genes of lung carcinoma are FGF2, CREB1, PPARG, RHOA,JUN,CXCL12,MET,CAV1,PIK3R1,SMAD4. Top 10 hub genes of lung carcinoma down-regulated genes are SERPINB8, SPP1, TAF9, AK6, PIK3CD, TEX10, GTF2E2,CD276,FLT1,PLCG1.We discovered the promising natural drug candidate effectively used in prognosis treatment against lung carcinoma, which are *Allium sativum, Arnebia euchroma* and *Nigella sativa.The* major compounds of these three drug candidate where identified and docked against the top 10 of both up regulated and down regulated hub genes.Allicin was proved to be the best indian medicinal drug target to be as a potential drug for lung carcinoma.Further studies on these natural drug targets are required in order to identify the proper and promising drug candidate alternatives that can be used to inhibit lung carcinoma.

## 7) APPENDIX

